# Therapeutic Communication Skills Training: An Effective Tool to Improve the Caring Behaviors of ICU Nurses

**DOI:** 10.1101/2020.02.12.945394

**Authors:** Farzane Zare, Jamileh Farokhzadian, Monirsadat Nematollahi, Sakineh Miri, Golnaz Foroughameri

## Abstract

**Background:** Caring behaviors are crucial in intensive care units (ICU) because patients in these wards require high levels of care. Effective communication with patients is one of the most important factors in caring behaviors of nurses working in ICUs. Therefore, the aim of this study was to evaluate the effect of therapeutic communication skills training on the caring behaviors of ICU nurese.

**Materials and Methods:** This experimental pre-test/post-test study was carried out on 105 nurses working in ICUs of hospitals affiliated to Yazd University of Medical Sciences in Iran in 2019. Nurses were randomly assigned into control (52 nurses) and intervention (53 nurses) groups. A two-day therapeutic communication training workshop was conducted for the participants. Data were collected using demographic information questionnaire and caring behaviors questionnaire before and one month after the intervention.

**Results:** The findings showed no significant difference between the control and intervention groups regarding the nurses’ caring behaviors in the pretest (P = 0.148). However, after implementation of the training program, a significant difference was observed in the mean scores of caring behaviors between the two groups. In the control group, the mean scores of caring behaviors decreased significantly after the intervention (P = 0.001); whereas, the mean scores of intervention group increased significantly after the intervention (P = 0.001).

**Conclusion:** According to the results, ICU nurses’ training in therapeutic communication skills had a positive effect on their caring behaviors. Therefore, we suggest the authorities prepare and implement educational packages of therapeutic communication skills as a coherent program for other nurses. As a result, the caring behaviors and the quality of cares can be improved for patients.

## Introduction

Intensive care unit (ICU) is one of the most important wards in hospitals. Due to the critical and complex conditions of patients, careful and standard provision of caring behaviors is crucial in these wards (1). The nurses of ICUs have more communication with the patients than the other health care providers; therefore, their caring behaviors cause a significant impact on the patients’ treatment and recovery (2). The important tasks of ICU nurses include conducting effective communication with patients, supporting patients, continuing the cares, and increasing the patients’ participation in treatment (3).

Caring behaviors in ICU consists of all critical cares necessary for the patients’ survival (4). These behaviors include the measures taken in relation to the well-being of patients, such as their comfort, sensitivities, and calmness by listening and paying attention to them, being honest to them, and accepting them without judgment. Consequently, the sense of security is enhanced in patients(5). The physical caring behaviors are the daily routines, diagnostic interventions, treatments, procedures, training, and problem solving. The psychosocial caring behaviors include trust, touch, body language, acceptance of emotions, faith, and honesty in behavior (6, 7). Shalaby et al. (2018) studied the caring behaviors of altruism, trust, hope, positive expression, education, and support in Jeddah hospitals and suggested that although ICU nurses emphasized the importance of these caring behaviors, they acted poorly in dealing with patients (6). Nurses’ caring behaviors increase the patients’ satisfaction and well-being, which consequently lead to the improved performance of the health care organizations (8).

Nurses require theraputic communication skills to show appropriate caring behaviors (9). Therapeutic communication refers to the process in which a nurse deliberately helps the patient’s improvement using verbal and non-verbal communications (10). In fact, it refers to a purposeful relationship between the patients and care givers. It is a tool for the therapist to communicate with the patient to create hope and positive change for the patient’s recovery(11). Admission in ICU creates anexiety for the patients (12) due to the disease, separation from family, immobility, and environmental noise, which can lead to patients’ impatience, depression, and irritability (13). In addition, ICU nurses have difficulty in communicating with patients using mechanical ventilation due to the lack of knowledge and skills. Therefore, they need education on communication skills that improves the patients’ care quality (9). Happ (2013) conducted a study in the USA over the effcet of intervention on nurse-patient relationship in ICU among the intubated, awake, and responsive patients. The findings showed that communicating with these patients was a common problem that caused distress and fear in patients and stress among nurses (14). A study reported that unconscious ICU patients were able to hear (15). Since hearing is the last sensation lost in patients with brain damage, speaking to and touching these patients were considered as important factors to communicate with these patients (16). A study in Canada investigated the effects of patient-based communication interventions on patients with communication disorders. The results showed that communicating with the patient had a positive effect on the patients’ recovery (17).

In general, the nurse-patient relationship improves the patient’s health. The care team should be aware that the patient-nurse relationship not only improves the patients’ disease, physical condition, and treatment, but also affects the patients’ physical, mental, and social health significantly (18). In fact, communication skills training can improve the care team ability to show their empathy and the patients’ ability to express their feelings (19). Popa-Velea et al. (2014) reported that the therapeutic communication training was necessary for the health care teams, especially physicians and nurses. Furthermore, they believed that communication barriers were tendency to judge, criticize, advise, and label patients, which lead to patients’ distrust. They stated that the type of words used to speak with the patients in order to transfer the sense of trust and empathy, the tone and melody of the voice, body language, honest attitude, and observance of the confidentiality principle were important in establishing a therapeutic relationship (20). Alasad et al. (2015) in Jordan argued that ICU patients wished to know more about the treatment measures taken for them. So, nurses should be trained about the importance of verbal and non-verbal communication with patients regardless of their prior knowledge. Nurses need knowledge and skills to create a therapeutic communication and are also required to learn the communication-based care (21). Therefore, due to the lack of study over the effect of communication skills on caring behavior of nurses in Iran, the present study was conducted. The aim was to determine the effect of communication skills training on caring behaviors of nurses who work in ICUs. The results of this study can be considered nurses’ therapeutic communication skills and caring behaviors.

## Methods and materials

### Design and Setting

This experimental pre-test/post-test study was conducted on the control and intervention groups. The study was carried out in ICUs (surgical and general ICUs, dialysis and cardiac ICUs) of hospitals affiliated to Yazd University of Medical Sciences in centre of Iran in 2019.

### Sample/participants

The study population included all nurses (N=300) working in the CCU, dialysis ward, surgical ICU, children ICU, and general ICU. The research participants were 110 nurses selected using the simple random sampling by the random number table. Then, they were categorized into control and intervention groups (55 nurses in each group). Finally, a total number of 52 members of the control group and 53 participants of the intervention group completed the study (N=105). The inclusion criteria consisted of having at least six months of work experience in ICU, Bachelor’s degree or higher in nursing, no mental or psychological diseases, no participation in the educational workshop of “therapeutic communication skills” during the past six months. The exclusion criteria included non-attendance in the educational program, lack of cooperation throughout the research, and incomplete questionnaire (22).

### Intervention procedure

The aims of this educational program were to familiarize the nurses with the therapeutic communication skills and to enhance their caring behaviors. In this study, a training program was carried out to promote the psychosocial and communication aspects of care in nurses. The two-day educational workshop was held for eight hours and included therapeutic communication skills (23) taught by the researcher, who was an MSc student in nursing, and a psychologist, who was a faculty member. In order to increase the nurses’ participation in the workshop, the intervention group was divided into two groups of 26 and 27 members and the workshops were held with the same training protocol in four days. The training curriculum was prepared and developed by the psychologist and researchers based on the literature review.

The educational content was approved by two faculty members in the field of psychiatric nursing. This content included interpersonal communication skills, communication goals, effective and ineffective communications, verbal and non-verbal communications and their barriers, effective listening, and empathy with the patient. In addition, the stages of communicating with patients, therapeutic communication techniques, communication barriers in nursing, skills to communicate with patients having mental problems such as anxiety, ability to communicate with patients having physical problems such as speech and audio disorders, and a brief education on body language (22-26). In this workshop, education was provided using educational slides, lectures, clips, questions and answers, brain storming, and practical trainings.

### Instruments

The data collection tool was a questionnaire consisting of two parts; the first was the demographic information questionnaire and the second part was the Larsson’s caring behaviors questionnaire. The demographic information questionnaire included the participants’ age, gender, marital status, educational level, employment status, service records in hospital and ICU, as well as service area. The standard caring behaviors questionnaire developed by Larson (1987) included 57 items dealing with six sub-skills of being accessible, explains and facilitates, comforts, anticipates, trusting relationships, monitors and follows through. The questions should be answered on a five-point Likert scale ranging from least important (1) to most important (5). The minimum score in this scale is 57, while the maximum attainable score is 285(27). The Persian version of this questionnaire was validated by Pashaei (2014) using the translation re-translation method. In order to determine the content validity, the experts’ view points were used and to determine the reliability, the test/re-test method was applied (r =0.87) (28).

### Outcome measurement

The caring behaviors’ questionnaire was used to measure the nurses’ care behaviors in control and intervention groups. The goal was to determine the effectiveness of the therapeutic communication skills on the improvement of caring behaviors in nurses. Prior to the intervention and one month after the intervention, questionnaires were distributed among the participants. During this period, the control group did not receive any training with regard to communication skills.

### Ethical consideration

The ethics Committee afliated to Kerman University of Medical Sciences approved this study with the code of (IR.KMU.REC.1396.1726). Initially, the hospital authorities were provided with an introduction letter and the necessary coordination was made to conduct the study. A cover letter explaining the study goals and the data collection procedure was also presented to the eligible participants before the data collection. Then, signed written consent forms were obtained from the participants and they were ensured about the data confidentiality and anonymity. Furthermore, they were expaliend about the voluntary participation in the study. Upon the completion of the intervention and collection of the second phase data, the therapeutic communication skills’ package was also administered to the control group in the form of a CD and handbook.

### Statistical analysis

Data were analyzed by SPSS version 21 using descriptive (frequency, percentage, mean, and standard deviation) and inferential statistics (independent samples t-test, paired t-test, and chi-square). The significance level was set at *P* < 0.05.

## Results

### Demographic characteristics

Most nurses in both groups were married women with a work experience of less than five years in ICUs. Most participants had a bachelor’s degree and were in the age range of 23-33. Table 1 shows the results of Chi-Square and Fisher’s exact tests. The findings showed that the participants of two groups were not significantly different in terms of demographic variables, except for their employment status.

**Table 1:**
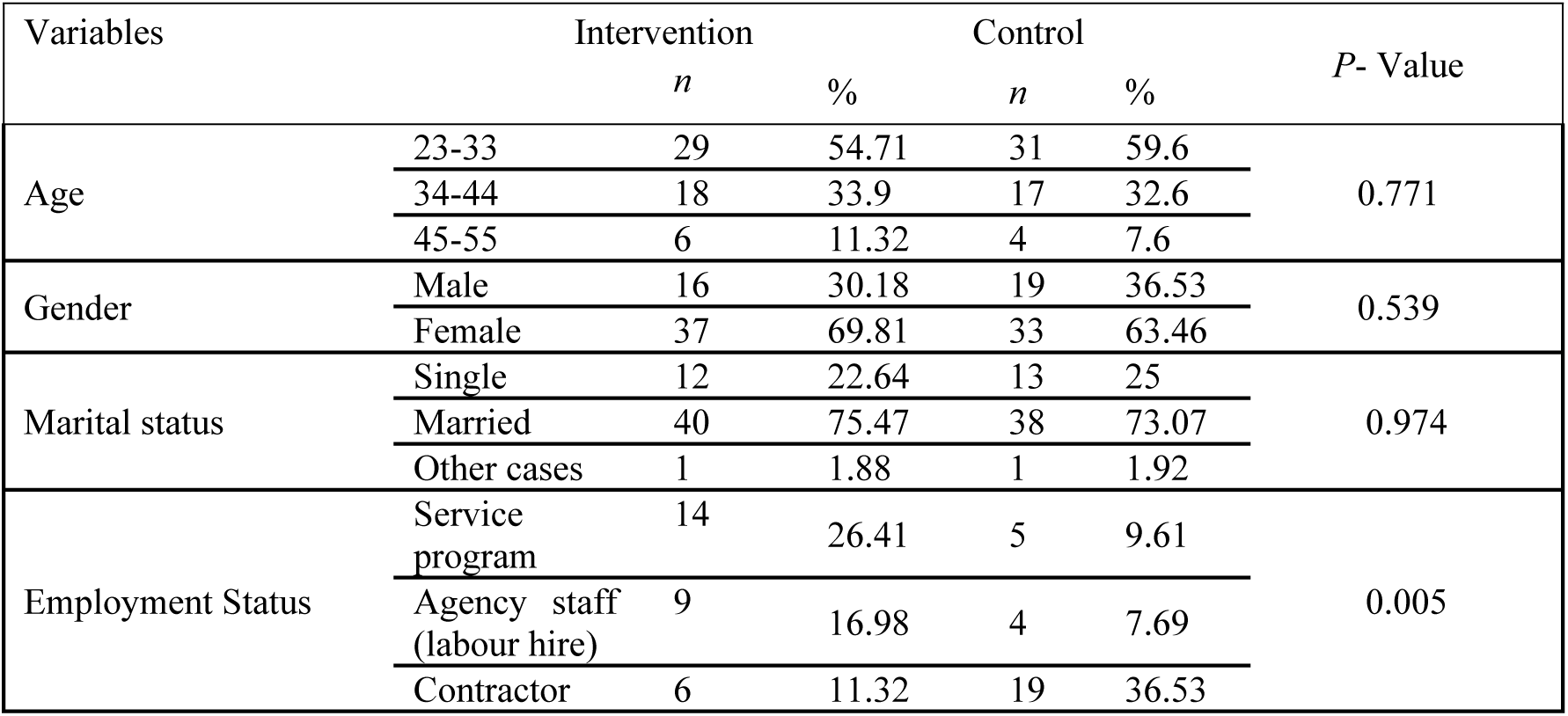
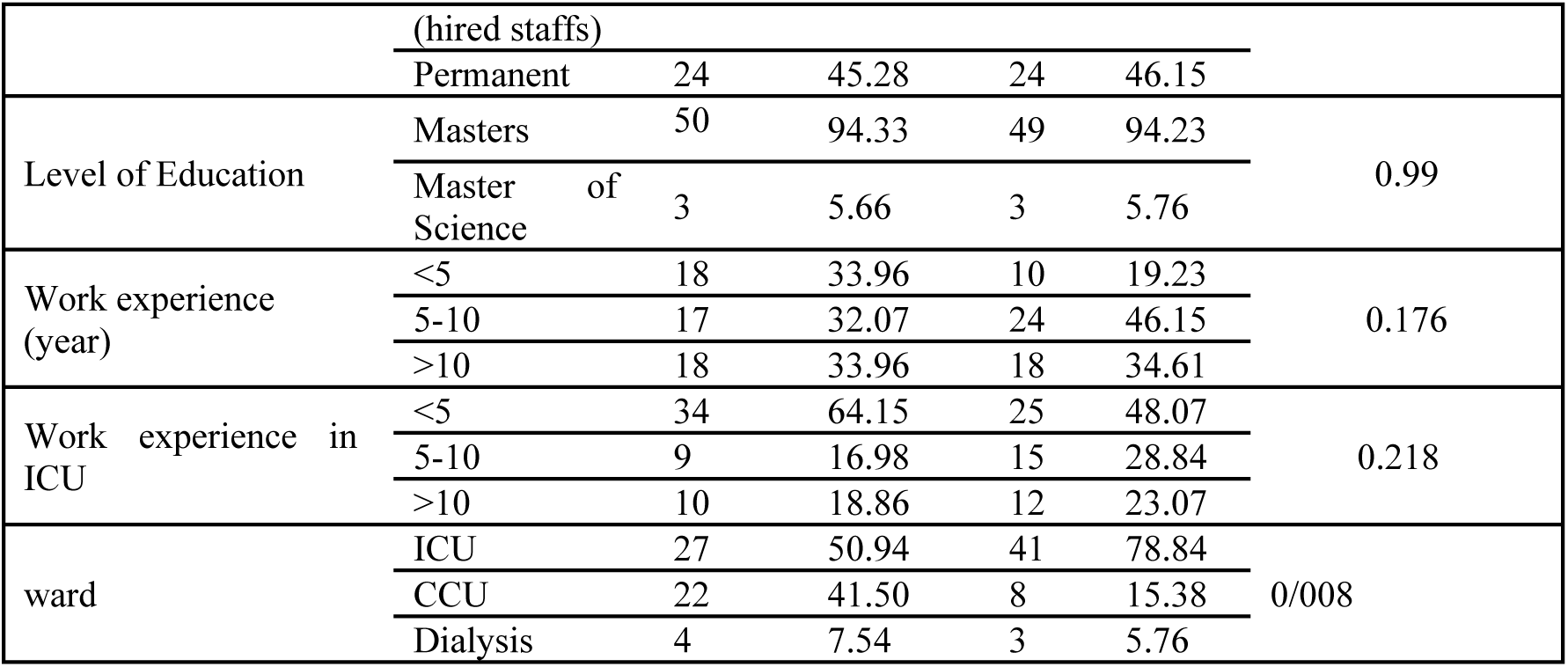
Comparison of demographic information of nurses in the intervention and control groups.

The results of Table 2 show that the mean score of caring behaviors in the intervention group (231.34 ± 18.17) was not significantly different from that of the control group (27.74 ± 238.28) (t = -1.45, P = 0.148). However, in the post-test stage, the mean score of caring behaviors in the intervention group (269.08 ± 4.92) increased significantly (t = 12.06, P = 0.001) in comparison with the control group (22.6.93 ± 68.22). Considering Table 2 that follows, post-test scores of the intervention group showed that the mean scores of all dimensions of caring behaviors improved significantly. In intervention group, of all the caring behavior dimensions, the highest mean difference was related to “trusting relationship” dimension (13.38), whereas, the lowest difference was related to “being accessible” dimension (2.97). This indicated that the educational program had the highest impact on the dimension of “trusting relationship” and the least impact on the dimension of “being accessible”. The paired t-test results indicated that after the education, caring behaviors and their dimensions improved significantly in the intervention group; whereas, they had a significant decrease in the control group (Table 2).

**Table 2:**
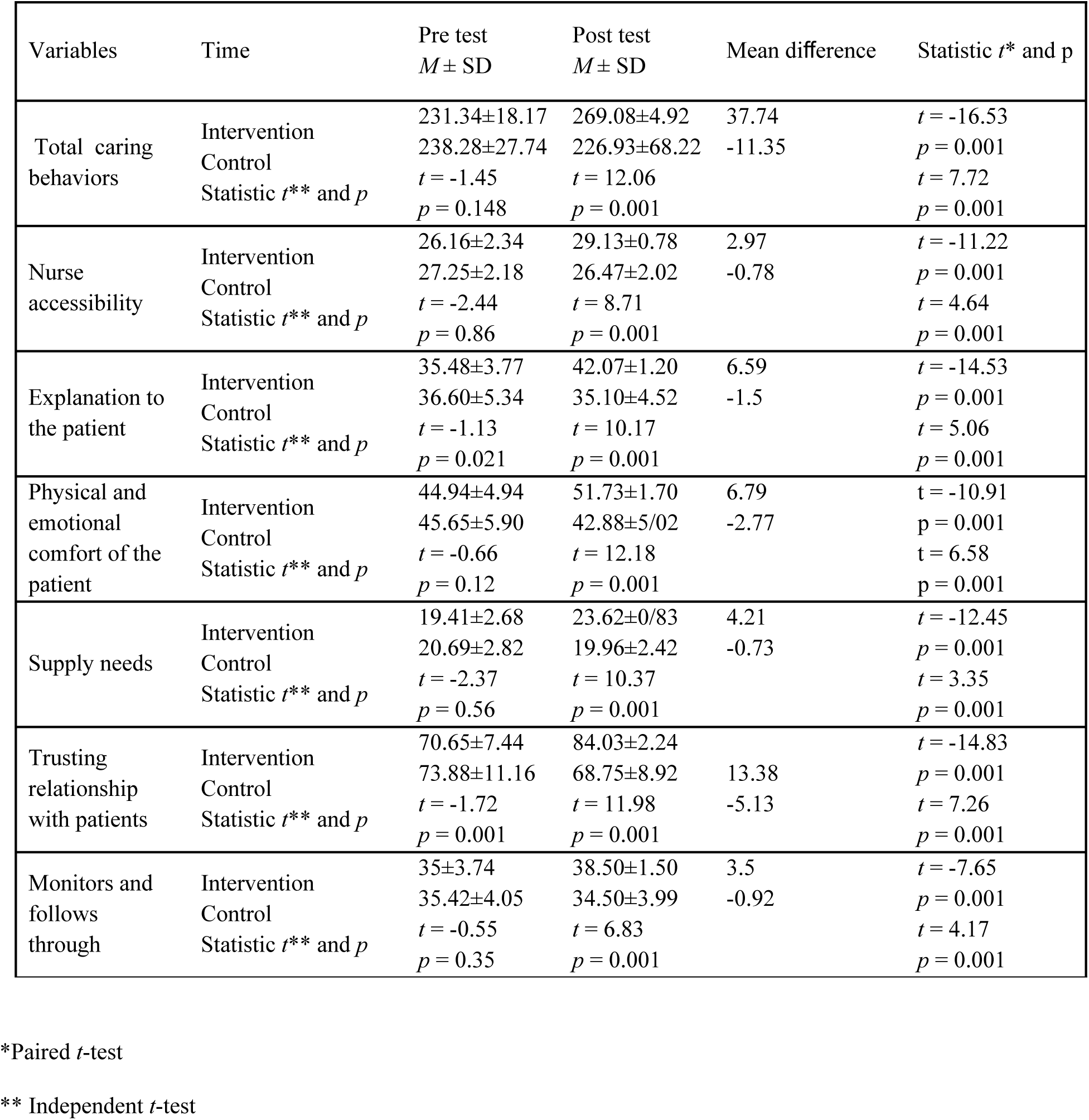
Comparison of mean scores of caring behaviors and its dimensions between the intervention and control groups at pre-test and post-test stage.

## Discussion

This study showed that the educational intervention improved the nurses’ caring behaviors in the intervention group. This indicates the positive effect of therapeutic communication skills on the caring behaviors of nurses. In a comprehensive review on the studies over the effect of communicative skills training on the caring behaviors of nurses, no similar study was found, which confirmes the novelty of our research. According to the results of numerous studies in Iran (33) and other countries (29-32), improving the communication skills enhanced the quality of nursing care. Porter et al. (2014) showed that nursing care behaviors were developed by applying the relationship-centered care professional model (34). Ratanawongsa et al. (2013) highlighted the importance of therapeutic communication skills in increasing the participation of patients in treatment, care, and acceptance of the diet (35).

Rask et al. (2009) evaluated the effect of communication skills training program on improving the nurse-patient communication. In contrast to our study, they reported that education did not have any significant effect on patients’ perception of nurses’ empathy, attention, and improved nursing care behaviors (36). It should be noted that the mentioned study did not address the nursing care behaviors. The discerpency between the results can be explained by the fact that in Rask’s study, patients were surveyed, but in the present study, the findings are based on the nurses’ self-reports.

The results showed that the highest mean difference of caring behaviors in the intervention group was related to the “trusting relationship” dimension, while the lowest difference was attributed to “being accessible” dimension. The “trusting relationship” dimension was considered as the most important factor in the current study because the educational content of therapeutic communication skills’ training as well as skills such as trust and empathy were emphasized in our study. In the same line with the present stusdy, the results of a study by Hillen et al. (2015) indicated that effective communication between the treatment staff and patients, especially application of non-verbal communication and body language led to a sense of trust in patients (37). The improvement of nurse-patient communication increased the patients’ trust and participation in treatment process and improved the quality of care (38). In the study by Pashaei et al. (2014), “trusting relationship” dimension had the highest priority for patients (28). The emphasis of accreditation standards of hospitals on the patients’ information confidentiality is another reason for the high priority of this dimension in the patients’ rights charter. In this regard, Berman et al. argued that observance of the information confidentiality is achieved by creating a nurse-patient trusted relationship (39). Movahedi et al. (2016) conducted a qualitative study in Iran and proposed that patients’ needs played an important role in the type of nurse-patient relationship in clinical settings. Therefore, the nurse-patient communication would increase by identification of the patients’ needs in different wards (40). The fact that ICU nurses take care of patients with higher needs might have led the participnats to consider the “trusting relationship” at the top priority.

Contrary to our results, some studies evaluated “trusting relationship” as a dimension with low priority. For example, Byrne et al. reported that nurses focused more on the patients’ physical care than psychological needs such as anxiety reduction in critical situations (41). Patients’ physical condition and prolonged hospitalization affect the nursing care in ICU and lead to prioritization of tasks related to the patients’ survival, while it may reduce the value of communicating with the patient (42). Variables such as patients’ gender, age, ethnicity, and health status, care providers and their underlying problems, the interpersonal skills, and organizational variables affect the development of trust between health care personnel and patients (43). This can justify the discerpencies in prioritization of different dimensions in various studies. Regarding these differences, other reasons can be the self-report method applied in the present study, evaluation of perceived caring behaviors without training communication skills in other studies, and prioritization based on a survey of patients and students (28, 44).

In addition, we found that education had the least effect on “being accessible” dimension, which can be explained by the nurses’ high work load, occupational burnout, and being neglected by high-level managers. These factors can prevent nurses from appropriate implementation of caring behaviors and effective communication with patients (45). The high workload of nurses, especially those in ICUs, low number of nurses, and lack of time make nurses just rely on the physical tasks and consider “being accessible” dimension less important (46). Contrary to our findings, a study in Iran (44) showed that “being accessible” and “patients’ monitoring and follow up” dimensions had the higest priority for students. Less important dimensions in this study were “explaining to patient”, “patient’s physical and emotional comfort”, “trusting relationship”, and “predicting the patients’ needs”. The students of this study selected “being accessible” dimension as the top priority, which indicates that they considered physical caring behaviors more important than the emotional behaviors and they were more concerned with the physical problems of patients. The priority of “being accessible” dimension and timely implementation of the therapeutic orders show the students’ most important caring behaviors that can be rooted in their education; their professors and teachers emphasized on and gave too much attention to the patients’ physical condition during the academic education.

## Limitations

This study has several limitations that should be considered by the furture researchers. One was the investigation of hospitals affiliated to Yazd University of Medical Sciences. The other constraint was lack of attention to the individual differences of participants. Moreover, the most important limitation of this study was the self-reporting data collection method. Therefore, we suggest application of observational checklists or patients’ surveys in the future studies. Another problem was the collection of information one month after the intervention. In order to achieve more accurate results, 3-6-month follow ups are recommended. Then, the results of different follow ups should be compared to determine the long-term impact of the training.

## Conclusion

The findings showed that therapeutic communication skills training significantly improved the caring behaviors in ICU nurses. This allows the nursing managers to use different educational approaches to enhance the ICU nursing care behaviors. Since communication skills training had a small effect on “being accessible” dimension, other interventions are required to improve it. Furthermore, considering the work difficulty and the type of patients in ICUs, the nursing managers should pay special attention to these nurses and provide better conditions for them to improve their caring behaviors. More qualitative studies are also suggested to evaluate the strategies for improving the nursing care behaviors in ICUs.

## Acknowledgments

The researchers appreciate all the nursing staff who generously offered their time to participate in the study.

